# DCGAN-Based Synthetic MRI Augmentation for Data-centric Brain Tumor Segmentation

**DOI:** 10.64898/2026.07.26.740722

**Authors:** Priyabrata Das, Jyotisman Rath, Amit Jaiswal, Bibhuti Bhusan Dash

## Abstract

Accurate brain tumor segmentation from magnetic resonance imaging (MRI) remains a challenging task because supervised deep learning models require large quantities of annotated data, which are expensive and time-consuming to obtain. This study investigates whether synthetic MRI images generated using a Deep Convolutional Generative Adversarial Network (DCGAN) can improve U-Net-based brain tumor segmentation using synthetic data augmentation.

Experiments were performed on the LGG-MRI dataset comprising 3,929 image-mask pairs. A baseline U-Net was first trained using the original training dataset. Synthetic MRI images were subsequently generated using a DCGAN, and threshold-derived pseudo-masks were assigned to the generated images to construct an augmented training dataset. The same U-Net architecture was then retrained using the augmented dataset and evaluated on an identical held-out test set. Compared with the baseline model, DCGAN-based augmentation increased the Dice coefficient from 0.2067 to 0.3037 and the Intersection over Union (IoU) from 0.1243 to 0.1918, while reducing the final test loss from 0.0474 to 0.0275. These results indicate that synthetic MRI augmentation was associated with improved segmentation performance under the experimental conditions of this study. However, the reliance on threshold-derived pseudo-labels and evaluation on a single dataset limit the generalizability of the results. The proposed workflow provides a reproducible implementation for evaluating DCGAN-based synthetic data augmentation in supervised brain tumor segmentation and establishes a baseline for future studies employing more reliable annotation strategies and broader experimental validation.

## 1. INTRODUCTION

Brain tumors exhibit substantial variation in their morphology, growth patterns, and anatomical location, making accurate segmentation from magnetic resonance imaging (MRI) inherently difficult [1], [2]. Tumor boundaries are often poorly defined because of heterogeneous tissue composition, infiltrative growth, necrotic regions, and the presence of surrounding edema [1], [2]. MRI remains the primary imaging modality for brain tumor evaluation because of its superior soft-tissue contrast and excellent visualization of intracranial anatomy [3]. Yet image acquisition is only one part of the problem. Delineating the tumor accurately still depends largely on manual annotation by experienced radiologists, particularly when lesion margins are diffuse or irregular [3], [4]. Although manual segmentation is regarded as the clinical reference standard, it is time-intensive, susceptible to inter-observer variability, and difficult to scale for the large annotated datasets required by supervised learning algorithms [4], [5]. These challenges have accelerated the development of automated brain tumor segmentation methods designed to improve consistency while reducing reliance on manual annotation [2], [5].

Convolutional neural networks (CNNs) have substantially improved the performance of automated medical image segmentation by learning hierarchical image features directly from training data [6], [7]. Among these architectures, U-Net remains the most widely adopted framework because its encoder–decoder architecture combines contextual information with fine spatial details, enabling accurate localization of anatomical structures [6], [8]. Numerous variants have subsequently been developed, incorporating residual connections, attention mechanisms, dense feature fusion, and, more recently, transformer-based modules to improve feature representation and segmentation accuracy [7], [9], [10]. Performance has improved steadily. The dependence on large, accurately annotated datasets has not. Regardless of architectural complexity, supervised segmentation models remain constrained by the quantity and quality of the available training data [7], [11]. This limitation is particularly pronounced in medical imaging, where generating pixel-level annotations requires substantial clinical expertise, time, and financial resources [11], [12].

Supervised segmentation models rely on the availability of large, accurately annotated datasets. In medical imaging, however, obtaining such datasets is rarely straightforward [11], [12]. Pixel-level annotation requires expert interpretation, substantial time, and consistent labeling across observers, making the process both expensive and difficult to scale [4], [11], [12]. Conventional data augmentation techniques, including image rotation, flipping, scaling, cropping, and elastic transformations, partially address this limitation by increasing the diversity of existing training samples [11], [12], [13]. Their ability to generate realistic anatomical variability, however, remains limited. Generative Adversarial Networks (GANs) offer an alternative by learning the underlying data distribution and synthesizing realistic images rather than simple transformed copies of existing samples [13], [14]. Consequently, GANs have been increasingly adopted for medical image augmentation, particularly in applications where annotated datasets are scarce [13], [14]. Nevertheless, generating realistic images alone does not solve the segmentation problem. Reliable annotations for synthetic images remain equally important, and the influence of different labeling strategies on downstream segmentation performance has not yet been fully established [13], [15].

Most existing studies evaluate generative models primarily in terms of image quality or visual realism, whereas their influence on downstream segmentation performance is often investigated using different datasets, network architectures, or experimental protocols [13], [15], [16]. Direct comparison between studies therefore remains difficult. Furthermore, synthetic images generated for supervised segmentation require corresponding annotations before they can be incorporated into model training. This step is frequently overlooked or addressed using application-specific labeling strategies, making it difficult to distinguish the contribution of synthetic image generation from that of the annotation approach [15], [16]. The present study adopts a deliberately simple and reproducible augmentation pipeline. A DCGAN was used to generate synthetic MR images, threshold-derived pseudo-masks were assigned to the generated samples, and the resulting dataset was used to retrain the same U-Net architecture under identical experimental conditions. This design isolates the effect of synthetic data augmentation without introducing additional architectural modifications or changes to the training strategy. Rather than proposing a new segmentation model, the study evaluates whether this augmentation workflow improves segmentation performance relative to a baseline model trained using the original dataset alone.

Motivated by these challenges, the present study evaluates the effect of DCGAN-based synthetic MR image augmentation on supervised brain tumor segmentation. A baseline U-Net model was first trained using the original LGG MRI Segmentation Dataset. Synthetic MR images generated by the DCGAN were subsequently incorporated into the training dataset after assigning threshold-derived pseudo-masks, and the same U-Net architecture was retrained under identical experimental conditions. Segmentation performance before and after augmentation was compared using the Dice coefficient, Intersection over Union (IoU), and test loss. The objective was not to develop a new segmentation architecture or modify the generative model. Instead, the study investigated whether a simple, reproducible augmentation pipeline could improve segmentation performance relative to a baseline model trained using the original dataset alone while providing a practical foundation for future investigations of GAN-assisted medical image augmentation.

## 2. METHODS

### 2.1 Dataset and Preprocessing

The experiments were conducted using the LGG MRI Segmentation Dataset, obtained from Kaggle [17]. The dataset comprised MRI scans from 110 patients, yielding 3,929 paired MR images and corresponding binary tumor masks for supervised brain tumor segmentation [17].

During dataset preparation, entries within the kaggle_3m directory were screened to identify valid patient folders containing paired MRI images and tumor masks. Non-patient entries, including README.md and data.csv, were excluded. Image files were retained only when the associated _mask.tif file was present, ensuring that each sample consisted of a correctly matched image–mask pair.

The complete dataset was partitioned into training and testing subsets using a 15% hold-out split with a fixed random seed (42), resulting in 3,339 training samples and 590 testing samples. A custom PyTorch Dataset class was implemented to load paired images and masks during model training and evaluation.

Each MR image was loaded in grayscale, resized to 256 × 256 pixels, and normalized to the range [0,1]. Corresponding tumor masks were resized to the same spatial resolution, converted to binary labels using an intensity threshold, and expanded to include a single-channel dimension. Images and masks were subsequently converted into PyTorch tensors and supplied to the network through DataLoader objects with a batch size of 16. Data shuffling was enabled for the training dataset and disabled for the testing dataset.

### 2.2 Baseline U-Net Segmentation

A custom implementation of the U-Net architecture was implemented as the baseline model for brain tumor segmentation [6]. The network accepted a single-channel 256 × 256 MR image as input and produced a corresponding single-channel probability map representing the predicted tumor region. The architecture followed the original encoder–decoder design comprising four encoding blocks, a bottleneck layer, and four decoding blocks connected through symmetric skip connections [6]. Following the original U-Net architecture, each convolutional block contained two consecutive 3 × 3 convolutional layers. In the present implementation, each convolution was followed by batch normalization and a rectified linear unit (ReLU) activation. Spatial downsampling within the encoder was performed using 2 × 2 max-pooling operations, whereas feature-map resolution in the decoder was restored using transposed convolution layers with a kernel size of 2 × 2. The bottleneck expanded the feature representation to 1024 channels before progressive reconstruction of the output segmentation map. A final 1 × 1 convolution followed by a sigmoid activation function generated pixel-wise probabilities for binary segmentation. The implemented network contained 31,042,369 trainable parameters.

Model training was performed using the Binary Cross-Entropy (BCE) loss function and the Adam optimizer with a learning rate of 1 × 10⁻³ and a weight decay of 1 × 10⁻⁵. The baseline model was trained for 15 epochs using the original training dataset and subsequently evaluated on the independent testing dataset before GAN-based data augmentation.

### 2.3 Deep Convolutional Generative Adversarial Network (DCGAN) Training

A Deep Convolutional Generative Adversarial Network (DCGAN) was implemented to generate synthetic MR images for data augmentation [18]. The network consisted of two adversarial components: a generator that synthesized realistic MR images from randomly sampled latent vectors and a discriminator that learned to distinguish generated images from real MR images [18].

The generator received a 100-dimensional latent vector sampled from a standard normal distribution and transformed it into a 256 × 256 single-channel MR image using a fully connected layer followed by four transposed convolutional layers. Batch normalization and rectified linear unit (ReLU) activation functions were applied after each intermediate transposed convolution, whereas a hyperbolic tangent (Tanh) activation function was used in the output layer. The generated images were subsequently rescaled to the intensity range [0,1] before further processing. The discriminator consisted of four convolutional layers with Leaky ReLU activation functions (negative slope = 0.2), batch normalization after intermediate convolutional layers, and a final sigmoid activation function that estimated the probability of an input image being real or synthetic.

To improve the representation of tumor morphology, the GAN was trained exclusively using MR images containing visible tumor regions. Training images were automatically selected from the original training dataset by retaining only slices whose corresponding tumor masks contained more than 1,000 foreground pixels, resulting in 1,141 tumor-containing MR images for GAN training. The generator and discriminator were optimized simultaneously using the Binary Cross-Entropy (BCE) loss function according to the adversarial objective

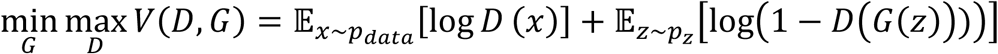

where *G* denotes the generator, *D* the discriminator, *x* a real MR image sampled from the training distribution *p_data_*, and *z* a latent vector sampled from the prior distribution *p_z_* [18].

Separate Adam optimizers were employed for the generator and discriminator using a learning rate of 2 × 10⁻⁴, *β*_1_ = 0.5, and *β*_2_ = 0.999. Training was performed for 20 epochs with a batch size of 16. Following completion of training, the generator was used to synthesize 3,339 MR images, corresponding to the number of samples in the original training dataset, for construction of the augmented training dataset.

### 2.4 Augmented Dataset Construction

Following DCGAN training, the generator was used to produce 3,339 synthetic MR images, corresponding to the number of samples in the original training dataset. Each generated image was saved individually for subsequent processing. Since pixel-wise annotations were unavailable for the synthesized images, binary pseudo-masks were generated by applying a fixed 8-bit grayscale intensity threshold of 100 to each image. Pixels with intensities greater than the threshold were assigned to the foreground class, whereas the remaining pixels were assigned to the background class. The generated images and their corresponding pseudo-masks were subsequently paired to form additional training samples.

The synthetic image-mask pairs were appended to the original training dataset without modifying the independent test dataset. Consequently, the number of training samples increased from 3,339 to 6,678, whereas the test dataset remained unchanged. An augmented PyTorch Dataset object and its corresponding DataLoader were then constructed using the same batch size (16) and data-loading configuration employed during baseline model training.

### 2.5 Augmented Model Training

To evaluate the effect of synthetic data augmentation, a new U-Net model with the same architecture was initialized and trained using the augmented training dataset containing both the original and DCGAN-generated image-mask pairs. No modifications were made to the network architecture, optimizer, loss function, learning rate, batch size, or training schedule. Consequently, the augmented model was trained under identical experimental conditions, with the composition of the training dataset representing the only experimental variable. Training was performed for 15 epochs using the augmented DataLoader, after which the trained model was evaluated on the unchanged independent test dataset. The trained network weights were subsequently saved for quantitative performance evaluation and comparison with the baseline model.

### 2.6 Performance Evaluation

Segmentation performance was evaluated using the Dice coefficient, Intersection over Union (IoU), and Binary Cross-Entropy (BCE) loss. Dice and IoU were computed on the independent test dataset after converting the predicted probability maps into binary segmentation masks using a decision threshold of 0.5. The BCE loss was calculated as

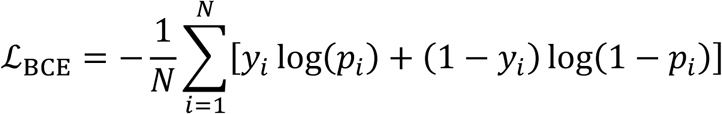

where *N* denotes the total number of pixels, *y_i_* represents the ground-truth label of pixel *i*, and *p_i_* denotes the predicted probability for the corresponding pixel.

The Dice coefficient quantified the spatial overlap between the predicted and reference tumor masks, whereas the IoU measured the proportion of the overlapping region relative to the combined segmented area [19], [20]. These metrics were computed as

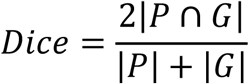

and

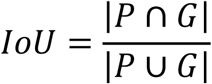

where *P* and *G* denote the predicted and reference segmentation masks, respectively, and *P* ∪ *G* represents their union [19], [20].

Mean Dice and IoU scores were calculated across all test samples for both the baseline and augmented models. In addition to the quantitative metrics, representative segmentation results were visualized by displaying the original MR image, the corresponding ground-truth mask, and the predicted segmentation. A direct comparison between the baseline and GAN-augmented models was subsequently performed using identical evaluation metrics and the same independent test dataset.

## 3. RESULTS

### 3.1 Baseline U-Net Segmentation Performance

The baseline U-Net model was trained using the original training dataset comprising 3,339 MR image-mask pairs for 15 epochs. Training and testing losses decreased rapidly during the initial epochs before gradually stabilizing as training progressed (Figure 2). The training loss decreased from 0.1562 to 0.0261, while the testing loss decreased from 0.0614 to 0.0474 over the 15 training epochs. A transient increase in the testing loss was observed during the later epochs; however, the overall trend remained stable, indicating stable convergence of the optimization process.

**Figure 1.**
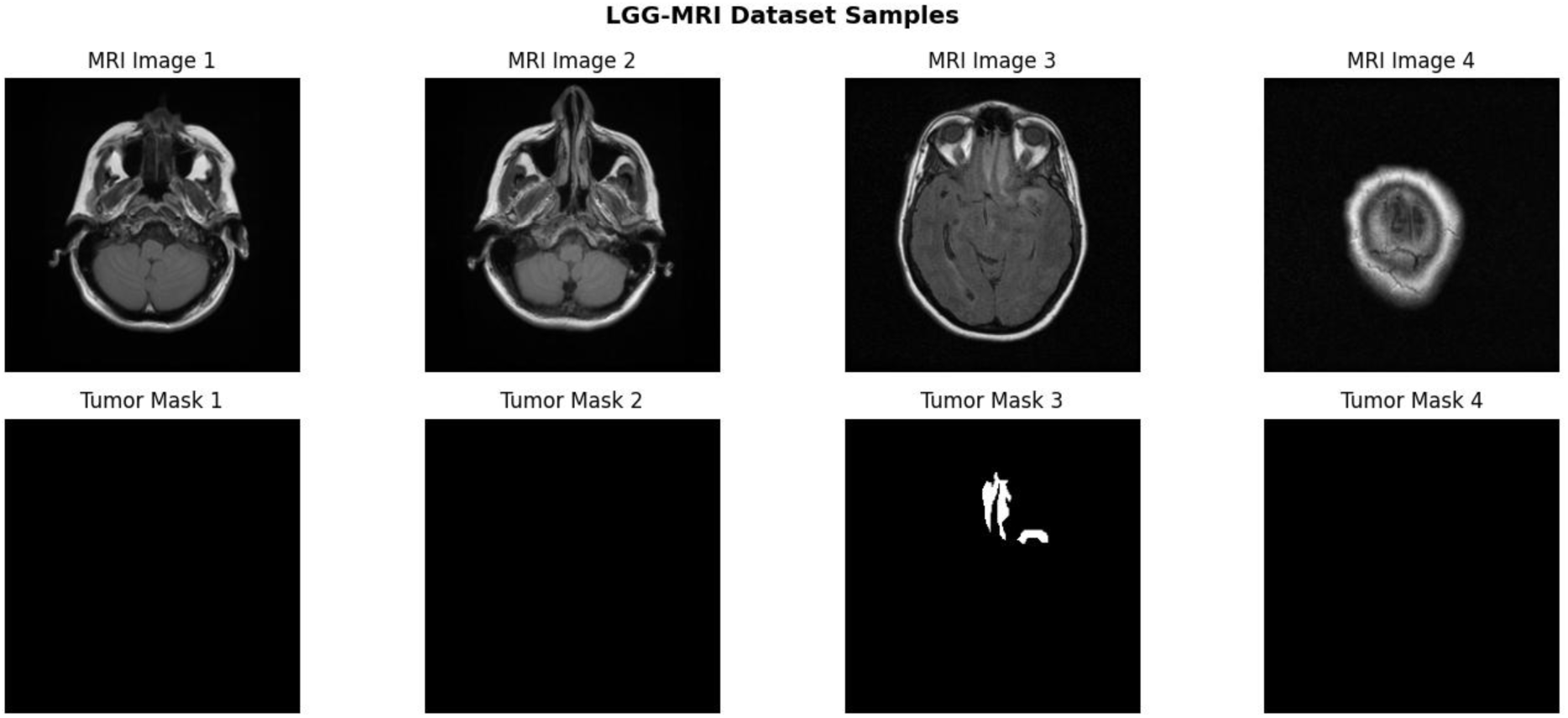
Representative MR images and corresponding binary tumor masks from the LGG-MRI Segmentation Dataset used for model development and evaluation.

**Figure 2.**
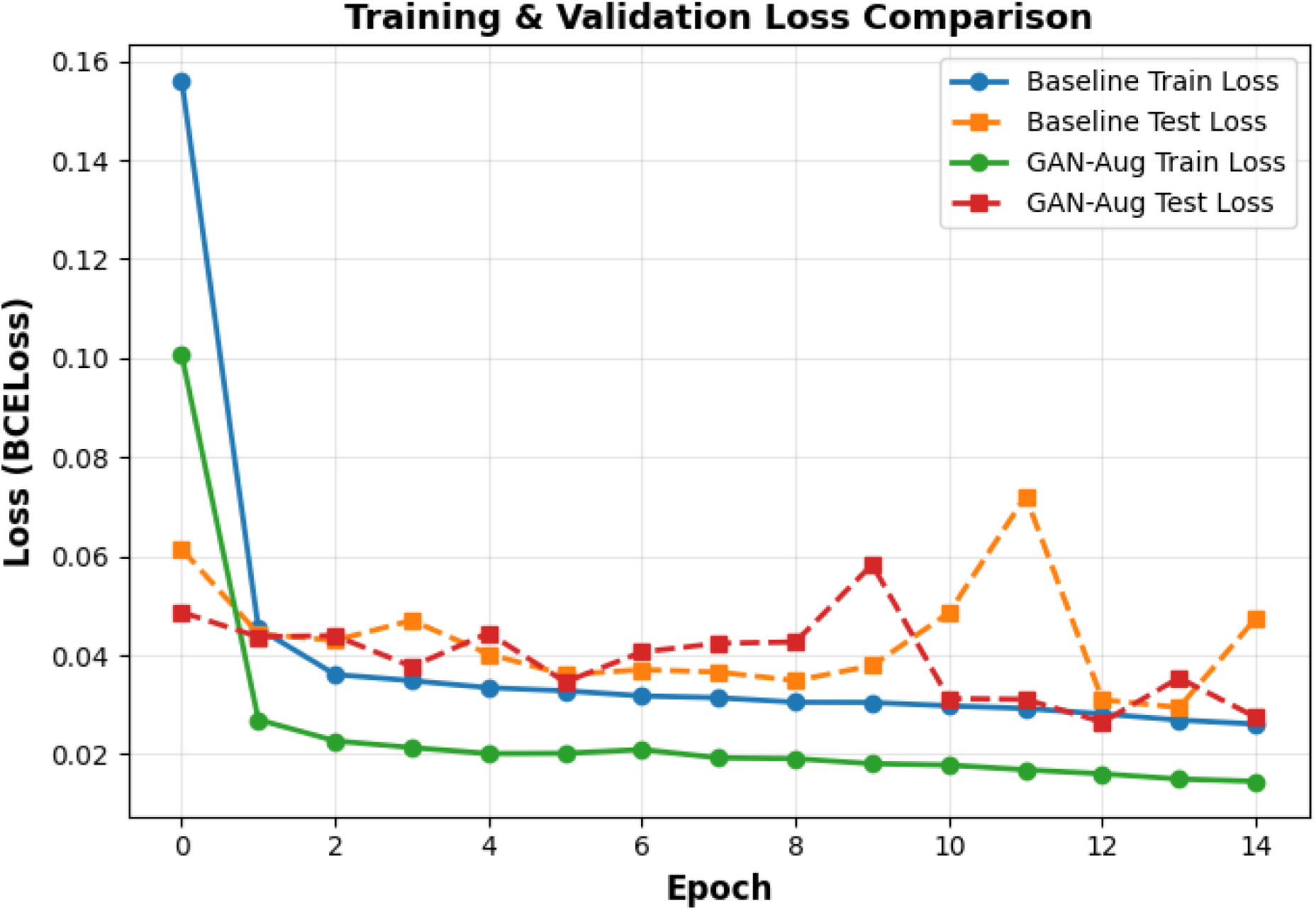
Training and testing Binary Cross-Entropy (BCE) loss curves for the baseline and GAN-augmented U-Net models over 15 training epochs.

The trained model was subsequently evaluated on the independent test dataset using the Dice coefficient and Intersection over Union (IoU). The baseline model achieved a mean Dice coefficient of 0.2067 and a mean IoU score of 0.1243. These values served as the baseline performance for subsequent comparison with the GAN-augmented model. These values served as the reference performance for evaluating the effect of GAN-based data augmentation in the subsequent experiments.

Representative predictions generated by the baseline model are presented in Figure 3. For each example, the original MRI image, manually annotated ground-truth mask, and corresponding predicted segmentation are displayed. The selected examples illustrate the qualitative segmentation performance of the baseline model across different test images. Since several displayed slices contain little or no visible tumor tissue, the quantitative evaluation metrics provide a more objective assessment of segmentation performance than the qualitative examples alone.

**Figure 3.**
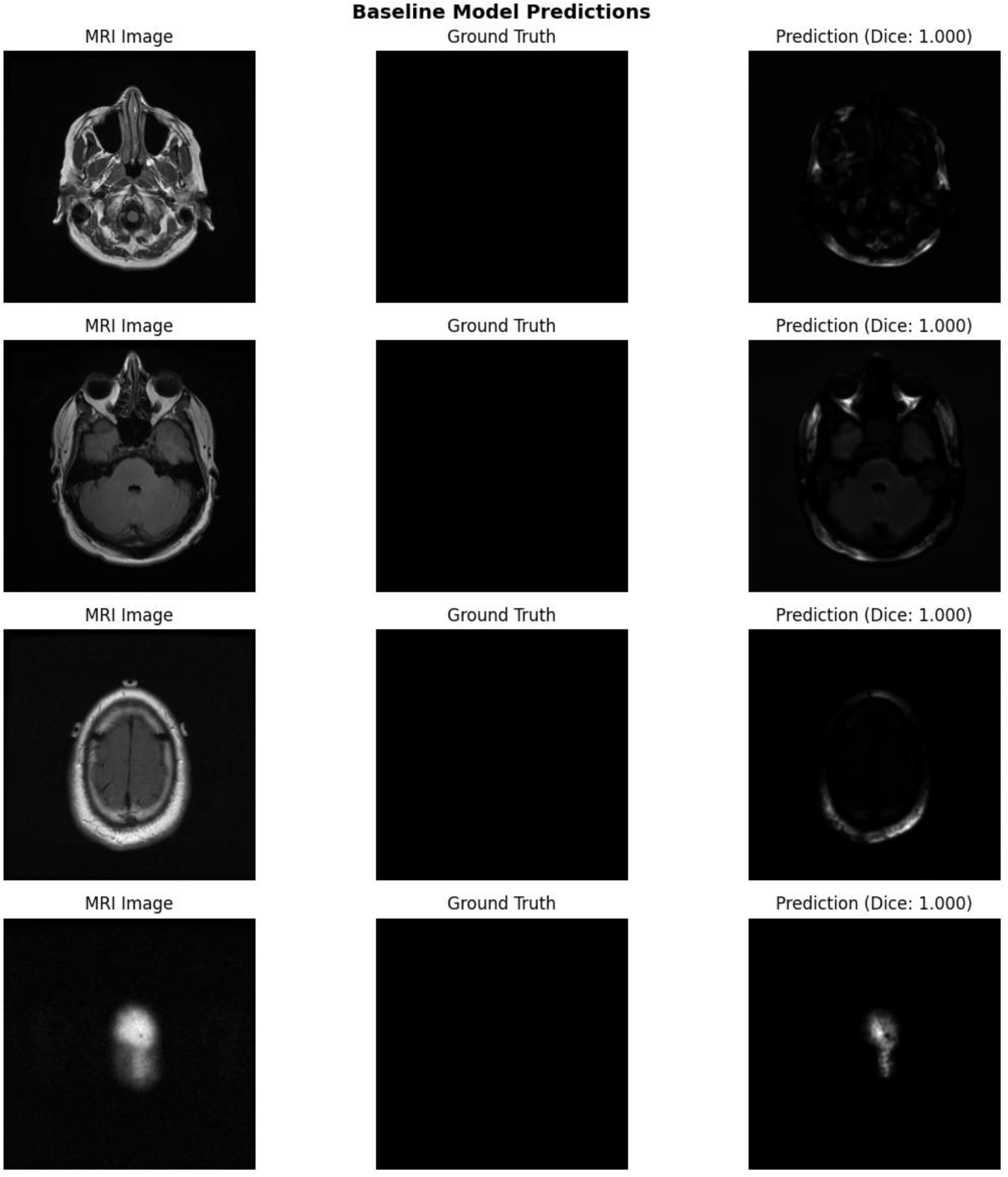
Representative segmentation results obtained using the baseline U-Net model. Each row shows the input MR image, corresponding ground-truth mask, and predicted segmentation.

### 3.2 DCGAN-Based Synthetic MRI Generation

To increase the diversity of the training data, a Deep Convolutional Generative Adversarial Network (DCGAN) [18] was trained using 1,141 MR slices containing visible tumor regions selected from the original training dataset. The adversarial training process was conducted for 20 epochs, during which the generator and discriminator were optimized simultaneously. The generator and discriminator loss curves are presented in Figure 4.

**Figure 4.**
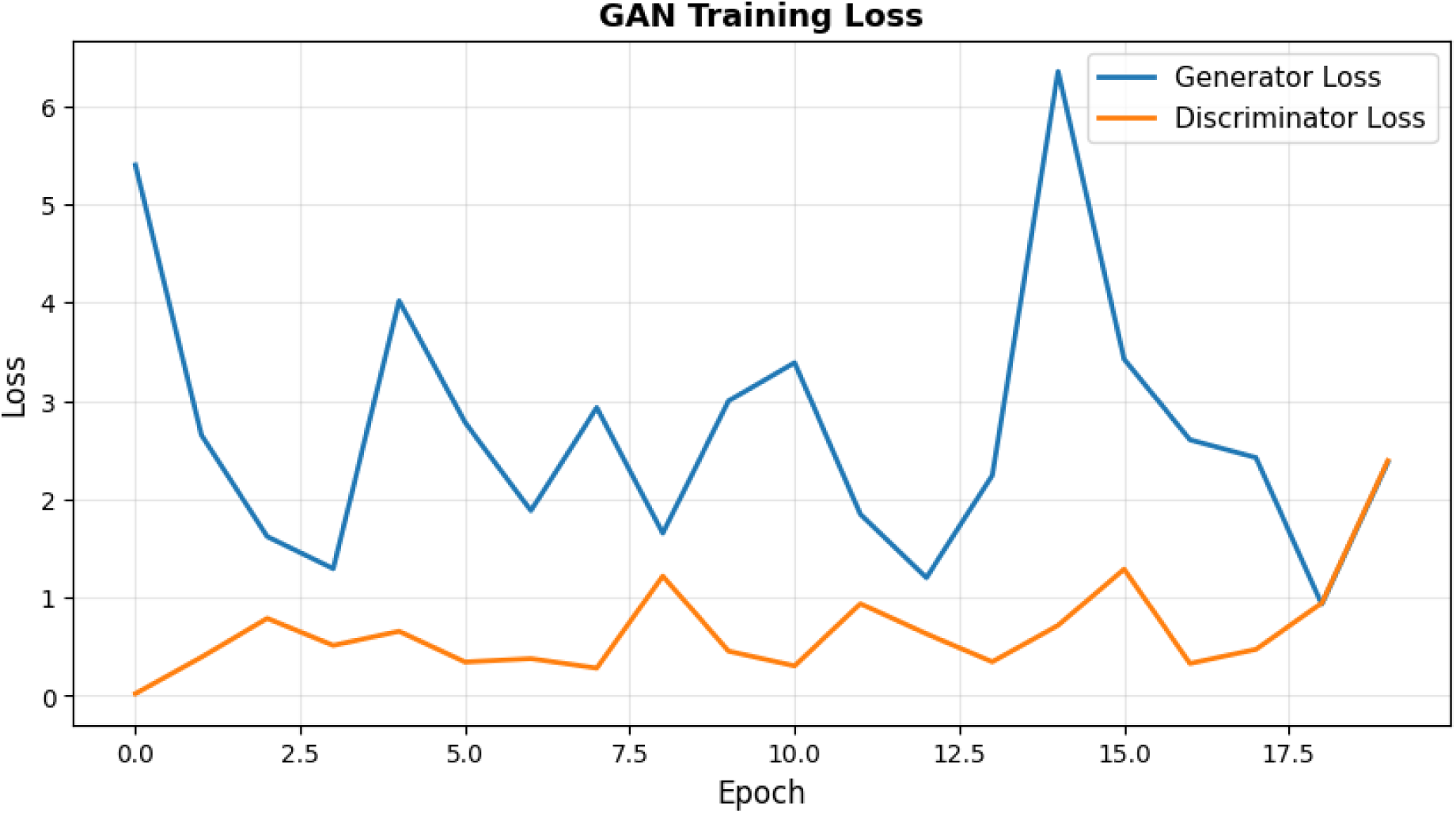
Generator and discriminator loss curves during DCGAN training over 20 epochs.

The loss profiles exhibited the oscillatory behaviour characteristic of adversarial optimization. During the early stages of training, both networks underwent rapid changes before gradually reaching a relatively stable equilibrium. Neither network consistently dominated the optimization process throughout training, indicating continued adversarial learning between the generator and discriminator.

Representative synthetic MR images generated by the trained DCGAN are shown in Figure 5. The generated images reproduced the overall anatomical structure of brain MR slices while displaying variations in tissue intensity and tumor appearance across different samples. Although minor structural artefacts remained visible in some generated images, the synthesized samples displayed sufficient anatomical variability for subsequent data augmentation.

**Figure 5.**
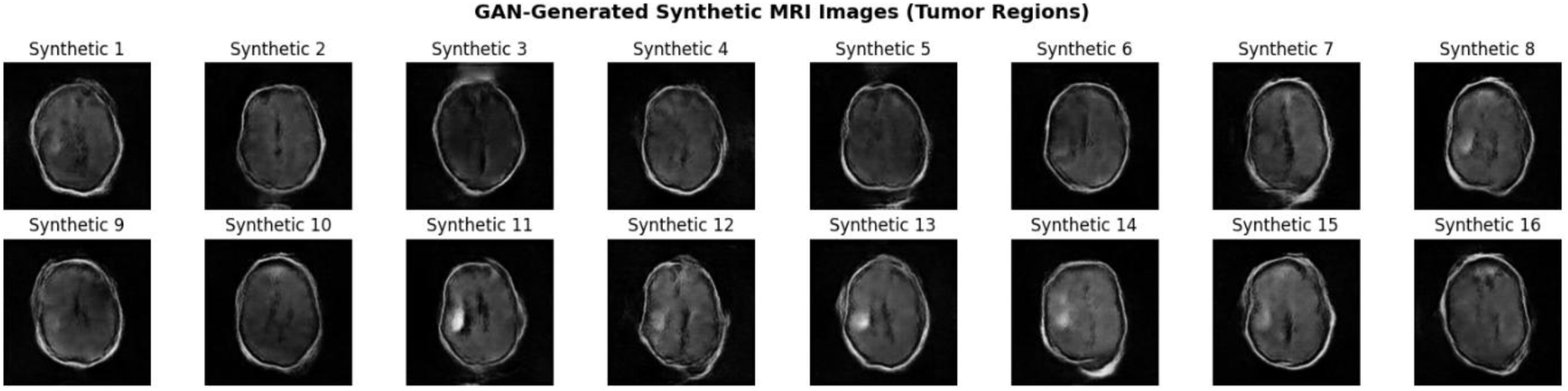
Representative synthetic MR images generated by the trained DCGAN following adversarial training.

### 3.3 Effect of GAN-Based Data Augmentation on Segmentation Performance

The synthetic MR image-mask pairs generated by the DCGAN were combined with the original training dataset to construct an augmented training set containing 6,678 MR image-mask pairs. A new U-Net model with the same architecture and training parameters as the baseline model was subsequently trained using the augmented dataset for 15 epochs.

Training and testing losses decreased progressively throughout optimization (Figure 6). Compared with the baseline model, consistently lower training and testing losses were observed during the later stages of optimization, the augmented model exhibited consistently lower training and testing losses during the later stages of training. At the final epoch, the training loss decreased to 0.0145, while the testing loss reached 0.0275, indicating improved convergence during model training on the augmented dataset.

**Figure 6.**
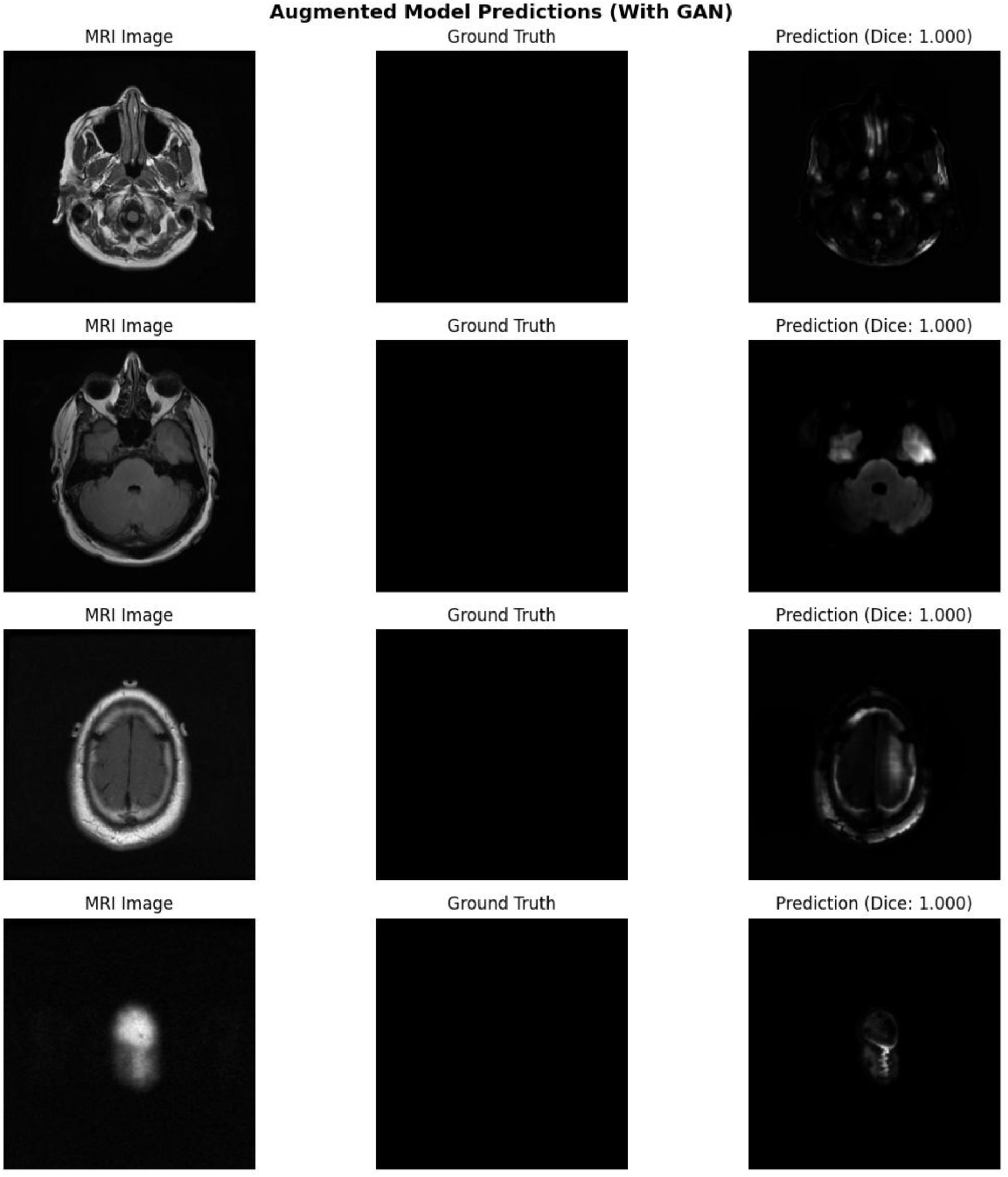
Representative segmentation results obtained using the U-Net model trained on the GAN-augmented dataset. Each row shows the input MR image, corresponding ground-truth mask, and predicted segmentation.

Evaluation on the independent test dataset demonstrated improved segmentation performance following GAN-based data augmentation. The augmented model achieved a mean Dice coefficient of 0.3037 and a mean Intersection over Union (IoU) score of 0.1918, compared with 0.2067 and 0.1243, respectively, for the baseline model. Representative segmentation results obtained using the augmented model are presented in Figure 7. The qualitative examples demonstrate the segmentation behaviour of the retrained network on the same test dataset used for baseline evaluation.

**Figure 7.**
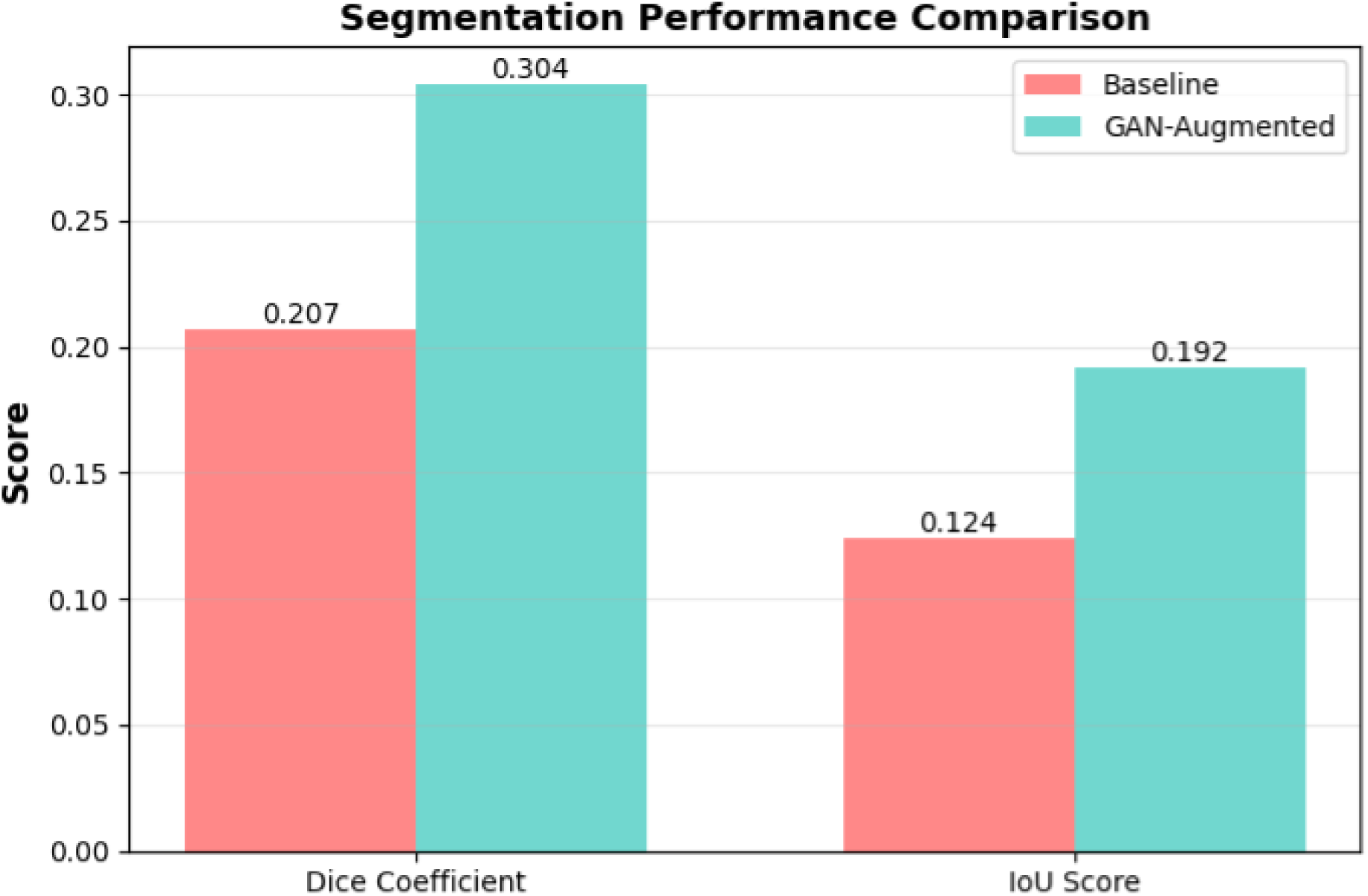
Comparison of the Dice coefficient and Intersection over Union (IoU) achieved by the baseline and GAN-augmented U-Net models.

### 3.4 Comparative Analysis of Baseline and GAN-Augmented Models

A direct comparison between the baseline and GAN-augmented U-Net models is presented in Figure 8 and Table 1. Training with the augmented dataset resulted in higher segmentation performance across all evaluated segmentation metrics. The mean Dice coefficient increased from 0.2067 to 0.3037, while the mean Intersection over Union (IoU) increased from 0.1243 to 0.1918. These correspond to relative improvements of 46.94% and 54.36%, respectively.

**Figure 8.**
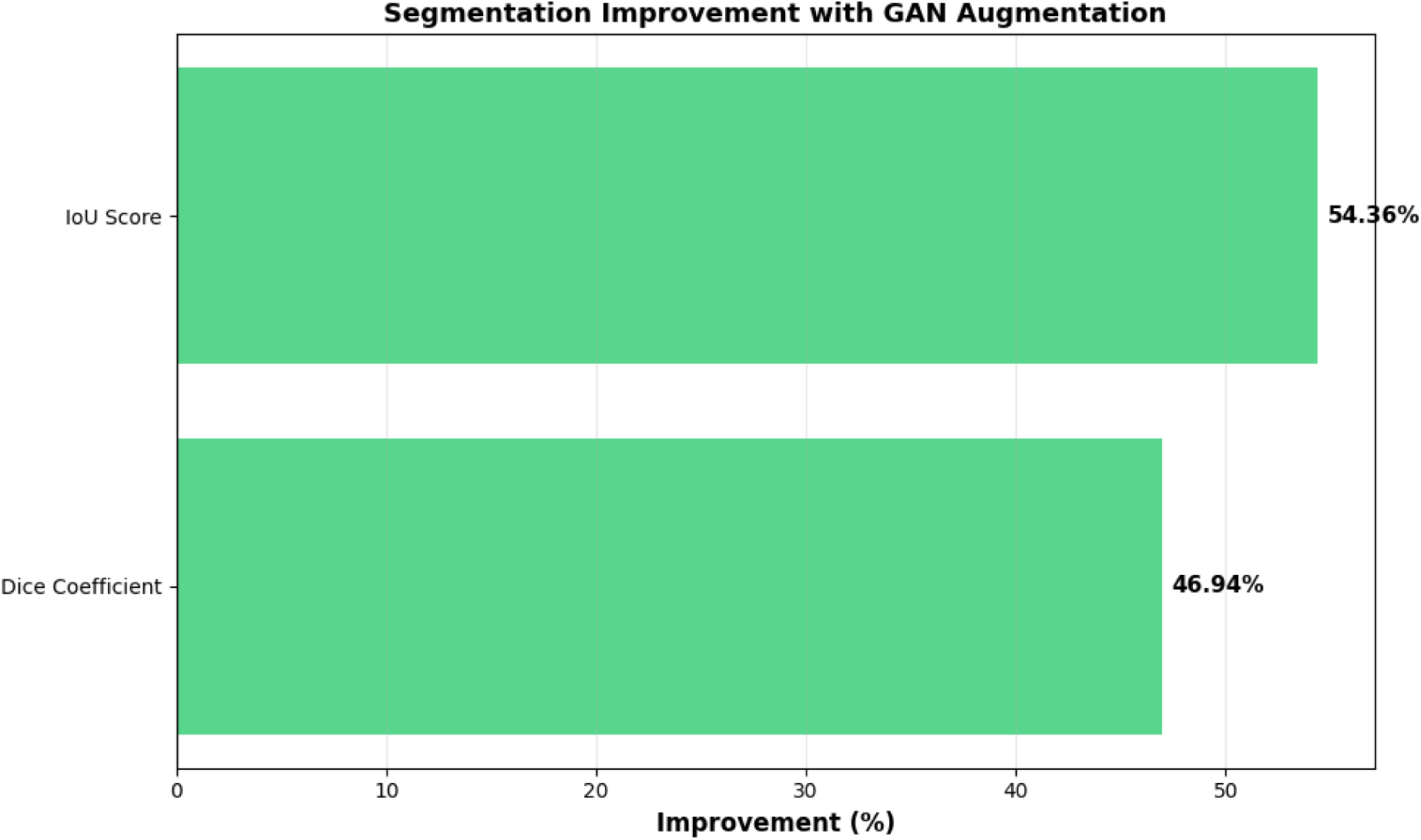
Relative percentage improvement in the Dice coefficient and Intersection over Union (IoU) following GAN-based data augmentation compared with the baseline model.

**Table 1.** Quantitative comparison of segmentation performance between the baseline and GAN-augmented U-Net models.

| Metric | Baseline U-Net | GAN-Augmented U-Net | Relative Improvement (%) |
| --- | --- | --- | --- |
| Dice coefficient | 0.2067 | 0.3037 | 46.94 |
| Intersection over Union (IoU) | 0.1243 | 0.1918 | 54.36 |
| Training dataset size | 3,339 | 6,678 | 100.00 |

The augmented model also exhibited lower training and testing losses during the final stages of optimization compared with the baseline model. These reductions in loss were accompanied by higher Dice and IoU scores on the independent test dataset. Since both models employed the same network architecture, optimizer, loss function, training schedule, and evaluation protocol, the observed differences in performance were attributed to the use of the augmented training dataset.

Overall, the quantitative metrics and qualitative segmentation results consistently demonstrated improved segmentation performance of the U-Net model following GAN-based data augmentation under the experimental conditions employed in this study.

## 4. DISCUSSIONS

### 4.1 Principal Findings

Brain tumor segmentation remains challenging because the performance of deep learning models depends heavily on the availability of annotated training data. In this study, incorporating DCGAN-generated synthetic MR images into the training dataset consistently improved the performance of the U-Net model. The augmented model achieved higher Dice and IoU scores while maintaining lower training and testing losses than the baseline model.

Importantly, the improvement was obtained without changing the network architecture or the optimization procedure. The only difference between the two experiments was the inclusion of synthetic training images. These findings indicate that expanding the training dataset through GAN-based augmentation was associated with improved segmentation performance on the independent test set. Although the absolute segmentation accuracy remained moderate, the results demonstrate that synthetic image generation can be a practical and reproducible strategy for improving model performance when annotated medical imaging data are limited.

### 4.2 Influence of GAN-Based Data Augmentation on Segmentation Performance

A major challenge in medical image segmentation is the limited availability of diverse, well-annotated datasets. Deep learning models trained on relatively small datasets are exposed to only a limited range of anatomical variations, tumor morphologies, and image characteristics during optimization. Consequently, their ability to generalize to previously unseen data may be compromised. Data augmentation has therefore become an important strategy for expanding the effective training distribution, and several recent studies have demonstrated the potential. of GAN-generated synthetic images to provide image variability beyond conventional augmentation techniques [11], [12], [13], [14], [15], [16].

The observations in the present study are consistent with these findings. The incorporation of GAN-generated MR images into the training dataset resulted in higher Dice coefficient and IoU values together with lower training and testing losses compared with the baseline model. Since the network architecture, optimizer, learning rate, loss function, training schedule, and evaluation protocol remained unchanged, the composition of the training dataset represented the primary experimental difference between the two models. Similar improvements following GAN-based augmentation have been reported for brain tumor segmentation and other medical image analysis tasks, supporting the observations of the present study [13], [14], [15], [16], [17].

An important aspect of the present work is the use of threshold-derived pseudo-masks for the generated synthetic images. Although these masks cannot provide the same level of precision as manually annotated ground-truth labels, the augmented model consistently achieved better quantitative performance than the baseline model on the independent test dataset. This observation suggests that carefully generated synthetic samples may provide useful complementary training information during model training even when perfectly annotated synthetic labels are unavailable. Nevertheless, pseudo-labels should be considered an augmentation strategy rather than a substitute for expert annotation. Future studies should investigate more advanced synthetic label generation methods together with rigorous validation using clinically annotated datasets [13], [17], [18], [19], [20].

### 4.3 Comparison with Previous Studies

The findings of the present study are broadly consistent with previous investigations demonstrating that GAN-based augmentation can improve the performance of deep learning models for medical image analysis, particularly when annotated datasets are limited [13], [14], [15], [16]. Several studies have reported that synthetic images generated using adversarial networks increase the diversity of the training data and reduce the risk of overfitting, leading to improved segmentation or classification performance across a range of medical imaging applications. The magnitude of improvement observed in the present study falls within the range reported for GAN-based augmentation approaches, although direct numerical comparison is complicated by differences in datasets, network architectures, and evaluation protocols. Although the magnitude of improvement varies depending on the imaging modality, network architecture, augmentation strategy, and dataset size, the overall trend reported in the literature supports the observations obtained in the present study.

Unlike many previous studies that employ manually annotated synthetic masks or more sophisticated label generation strategies, the present work adopted a comparatively simple threshold-based pseudo-labeling approach for the generated images. Despite this simplified methodology, the augmented model achieved higher Dice coefficient and IoU values than the baseline model. This suggests that improvements in segmentation performance can still be achieved using relatively straightforward augmentation pipelines, although the quality of the generated labels is expected to influence the final model performance.

Another distinction of the present study is the use of a conventional U-Net architecture without modifications to the network design. Recent brain tumor segmentation studies have increasingly focused on attention mechanisms, transformer-based architectures, and hybrid deep learning models [7], [8], [9], [10]. In contrast, the objective of the present work was not to develop a new segmentation architecture but to evaluate whether improving the diversity of the training data alone could enhance model performance. The observed improvements indicate that data augmentation remains an important component of segmentation workflows, independent of improvements in network architecture.

### 4.4 Study Limitations

Several limitations of the present study should be considered when interpreting the findings. First, the experiments were performed using a single publicly available low-grade glioma MRI dataset. Although this dataset is widely used for benchmarking brain tumor segmentation algorithms, evaluation on additional datasets acquired from different institutions and imaging protocols would provide a more comprehensive assessment of model robustness and generalizability.

Second, the synthetic masks associated with the generated MR images were obtained using a threshold-based pseudo-labeling strategy rather than manual expert annotation. While this approach enabled efficient construction of an augmented training dataset, it may not accurately represent complex tumor boundaries. Consequently, the quality of the synthetic labels is likely to influence the performance of the segmentation model.

Another limitation is that the quality of the generated synthetic images was assessed qualitatively through visual inspection. Quantitative measures of image realism, such as Fréchet Inception Distance (FID), Structural Similarity Index Measure (SSIM), or expert radiological evaluation, were not included in the present study. Incorporating these assessments would provide a more comprehensive evaluation of the generated data. Each model configuration was trained and evaluated once. Consequently, statistical variability arising from repeated training runs with different random initializations was not assessed.

Finally, the study focused exclusively on evaluating the effect of GAN-based data augmentation using a conventional U-Net architecture. More recent segmentation networks, including attention-based and transformer-based models, were not investigated. Whether similar improvements can be achieved across different architectures remains an important direction for future research.

### 4.5 Future Perspectives

The findings of the present study provide several directions for future research. One immediate extension would be to replace the threshold-based pseudo-labeling strategy with synthetic masks generated using dedicated segmentation networks or expert-annotated labels. Improving the quality of the synthetic labels may further enhance the effectiveness of GAN-based data augmentation.

Future studies should also evaluate the proposed augmentation strategy using larger and more diverse multi-institutional datasets to determine its robustness across different imaging protocols and patient populations. Such validation would provide a more comprehensive assessment of the generalizability of the approach.

Although the present study focused on a conventional U-Net architecture, the augmentation strategy is independent of the segmentation network. Evaluating GAN-generated synthetic images with more recent architectures, including attention-based, transformer-based, and foundation-model-based segmentation architectures, may provide further insight into the broader applicability of synthetic data augmentation.

Finally, quantitative evaluation of the generated images using dedicated image quality metrics together with external clinical validation would provide a more rigorous assessment of the usefulness of synthetic medical images in brain tumor segmentation. Such investigations may help establish standardized evaluation frameworks for the integration of generative models into medical image analysis pipelines.

## 5. CONCLUSION

This study investigated the effect of Deep Convolutional Generative Adversarial Network (DCGAN)-based synthetic MR image augmentation on brain tumor segmentation using a U-Net architecture. A baseline segmentation model trained on the original LGG-MRI dataset was compared with an identical model trained using an augmented dataset containing both original and GAN-generated synthetic images. Under identical training conditions, the GAN-augmented model achieved higher Dice coefficient and Intersection over Union (IoU) values together with lower training and testing losses than the baseline model.

The findings demonstrate that GAN-based synthetic image augmentation can improve segmentation performance without requiring modifications to the underlying network architecture or training strategy. The proposed workflow provides a simple and reproducible baseline for investigating the contribution of GAN-generated synthetic images to supervised brain tumor segmentation. Although the study employed threshold-derived pseudo-labels and was evaluated using a single publicly available dataset, the results suggest that synthetic image generation represents a practical and reproducible strategy for increasing training data diversity when annotated medical images are limited. Future studies should evaluate the proposed workflow using larger multi-institutional datasets, improved synthetic label generation methods, and contemporary segmentation architectures to determine its broader applicability in brain tumor segmentation.

